# Viral Airway Injury Promotes Cell Engraftment in an *In Vitro* Model of Cystic Fibrosis Cell Therapy

**DOI:** 10.1101/2022.11.14.516213

**Authors:** Rhianna E. Lee, Teresa M. Mascenik, Sidra C. Major, Catherine A. Lewis, James E. Bear, Raymond J. Pickles, Scott H. Randell

**Affiliations:** Marsico Lung Institute/Cystic Fibrosis Research Center, University of North Carolina at Chapel Hill, NC 27599; Department of Cell Biology and Physiology, University of North Carolina at Chapel Hill, NC 27599; Department of Microbiology and Immunology, University of North Carolina at Chapel Hill, NC 27599

**Keywords:** Cell therapy, cell engraftment, airway epithelium, competition, radiation, viral injury

## Abstract

Cell therapy is a potential treatment for cystic fibrosis (CF). However, cell engraftment into the airway epithelium is challenging. Here, we model cell engraftment *in vitro* using the air-liquid interface (ALI) culture system by injuring well-differentiated CF ALI cultures and delivering non-CF cells at the time of peak injury. Engraftment efficiency was quantified by measuring chimerism by droplet digital PCR and functional ion transport in Ussing chambers. Using this model, we found that human bronchial epithelial cells (HBECs) engraft more efficiently when they are cultured by conditionally reprogrammed cell (CRC) culture methods. Cell engraftment into the airway epithelium requires airway injury, but the extent of injury needed is unknown. We compared three injury models and determined that severe injury with partial epithelial denudation facilitates long-term cell engraftment and functional CFTR recovery up to 20% of wildtype function. The airway epithelium promptly regenerates in response to injury, creating competition for space and posing a barrier to effective engraftment. We examined competition dynamics by time-lapse confocal imaging and found that delivered cells accelerate airway regeneration by incorporating into the epithelium. Irradiating the repairing epithelium granted engrafting cells a competitive advantage by diminishing resident stem cell proliferation. Intentionally causing severe injury to the lungs of people with CF would be dangerous. However, naturally occurring events like viral infection can induce similar epithelial damage with patches of denuded epithelium. We found that viral preconditioning promoted effective engraftment of cells primed for viral resistance.

## Introduction

Cystic fibrosis (CF) is a chronic genetic disease caused by mutations in the *cystic fibrosis transmembrane regulator (CFTR)* gene, leading to absent or non-functional CFTR protein. In the lungs, diminished CFTR leads to defective chloride and bicarbonate transport, increased mucus viscosity, impaired mucociliary clearance, and chronic bacterial infection and inflammation (1). Though CF is monogenic, there are over 2000 *CFTR* variants (2) with effects ranging from no protein synthesis (class I), defective CFTR trafficking (class II), reduced channel conductance (class III), partial CFTR function (class IV), deficient protein synthesis (class V), and shortened protein half-life (class VI). Small molecule CFTR modulators improve CFTR trafficking and gating for individuals with class II and III variants (3–7). However, these treatments are *CFTR* variant-specific and unlikely to help people with low *CFTR* mRNA or protein levels.

CF has long been considered an ideal candidate for gene therapy, but clinical trials to date have been minimally successful (8, 9). This is largely due to gene delivery barriers including airway mucus, epithelial tight junctions, and the organization of the airway epithelium itself. Access to the basal stem cell compartment is blocked by differentiated columnar cells. Thus, gene therapy effects are short-lived, lasting only until the corrected cells turnover. Autologous cell therapy is another potential approach to treatment. Theoretically, airway epithelial cells could be collected by bronchiolar lavage, from sputum, or via bronchoscopy, expanded and gene-corrected *ex vivo,* and delivered to the stem cell niche where corrected cells would engraft, differentiate, and ultimately correct disease phenotypes. However, cell engraftment in the airways has historically proven challenging. Previous work has shown that effective cell engraftment will require epithelial disruption to create space for delivered cells (10–14), but the extent of airway preconditioning required is unknown.

Many questions remain about the optimal cell therapy approach, including: 1) does the cell expansion method impact the “engraftability” of delivered cells; 2) what are the minimum preconditioning requirements for long-term cell engraftment; 3) how do the delivered cells and the repairing epithelium interact; and finally, 4) will engrafted cells differentiate and restore CFTR function? To answer these questions, we developed an *in vitro* engraftment model using the air-liquid interface (ALI) culture system to enable systematic comparisons of delivered cell populations and preconditioning regimens. Molecular biology techniques, electrophysiology assays, and high-resolution imaging are well-established in the ALI culture system, permitting in-depth study of cell engraftment dynamics and outcomes.

With the ALI model, we found that severe injury characterized by partial epithelial denudation is required to promote effective and persistent cell engraftment in the airway epithelium. Injured airway cells repaired rapidly and competed with delivered cells for space. Though the injury requirements are high, the payoff of cell engraftment was substantial with delivered cells constituting 28% of the final culture and rescuing 20% of wildtype CFTR function. Intentionally causing severe injury to the lungs of people with CF would be dangerous and translationally unrealistic. However, naturally occurring events like viral infection can induce similar epithelial damage with luminal cell shedding and patches of denuded epithelium (15, 16). We found that viral preconditioning promoted effective engraftment of cells primed for viral resistance *in vitro.* This finding may abrogate the need for intentional airway injury and allow us to take advantage of naturally occurring injury events to promote cell engraftment for CF cell therapy. Preliminary versions of this report have been published in abstract form (17–19).

## Materials and Methods

### Cell culture

Human bronchial epithelial cells (HBECs) from CF and non-CF donors were expanded as described (20) before culturing at an air-liquid interface (ALI) for 21-60 days (d) in 6.5-mm Transwell (Corning 3470) or 12-mm Millicell (MilliporeSigma PICM01250) inserts. Non-CF HBECs were expanded to passage 1 (P1) by conventional (21) or conditionally reprogrammed cell (CRC) culture methods (22) described previously. Cells were transduced with an GFP-expressing adenovirus with 10 μg/mL polybrene for 24 hours (24h) where indicated.

### Engraftment

Well-differentiated CF ALI cultures were washed with PBS for 20 minutes (20 min) and treated 24h later with PBS (ThermoFisher J61196-AP), 30 mM sodium caprate (C10; MP Biomedicals ICN221694), or 0.125% or 0.25% polidocanol (PDOC; Sigma P9641) for 5 min. Non-CF cells were added apically 5 min after PBS or C10 treatment or 24h after PDOC treatment by adding 6 × 10^4^ cells suspended in 100 μL of ALI differentiation media for Transwell inserts or 2 × 10^5^ cells suspended in ALI differentiation media for Millicell inserts (Supplemental Figure 1). Cell cultures were irradiated with 2 gray (Gy) radiation 24h after PDOC injury and immediately before cell addition where indicated. For viral preconditioning, CF ALI cultures were inoculated with 100 μL of influenza-A (1.5 × 10^7^ PFU/mL) or mock (ALI media) for 2h. After 5d, cultures were treated with vehicle (ALI media) or ribavirin (100 μg/mL) for 72h. Non-CF cells were treated ± 1.2 ng/mL IFN-λ for 24h and engrafted onto mock- or influenza-treated CF cultures at 6d post-inoculation (Supplemental Figure 2).

### Analysis

Transepithelial electrical resistance (TEER) was measured using an EVOM2 device. Droplet digital PCR (ddPCR) was performed by probing for *AMELX* and *AMELY* genes in male-female mis-matched cultures as previously (23). Ussing measurements were performed 14d after engraftment using amiloride (Amil), forskolin (FSK), CFTR inhibitor-172 (CFTRinh-172), and uridine-5’-triphosphate (UTP) as previously described (24). Time-lapse confocal imaging of engrafted ALI cultures stained with Calcein green (Invitrogen C34852) or Calcein red-orange (Invitrogen C34851) was performed using a Zeiss 880 Airyscan. Cultures were imaged immediately after cell addition every 20-25 min for 20h. The repairing epithelium was manually traced and the area was quantified in imageJ.

### Whole mount immunostaining

Cultures were fixed in 4% paraformaldehyde and stained with antibodies against α-tubulin (Millipore MAB1864; 3 μg/mL), MUC5AC (ThermoScientific 45M1; 4 μg/mL), and E-cadherin (Cell signaling Technology 24E10; 3 mg/mL) and counterstained with Hoechst (Invitrogen 33342) and phalloidin (Invitrogen A22287) as previously described (25). The Click-iT EdU kit (Invitrogen C10337) was used to visualize proliferating cells. Cultures were imaged using a Leica TCS SP8 (40X objective) or an Olympus VS200 slide scanner (10X objective). Nuclei were quantified using imageJ. Ciliated cell count and MUC5AC signal was obtained using Volocity (PerkinElmer). The denuded membrane area was measured using Adobe Photoshop.

### Statistics

Statistics were performed using GraphPad Prism 9. Data is reported as the mean ± standard error of the mean (SEM). Unpaired T-test, one-way analysis of variance (ANOVA) with Tukey post-test, and two-way ANOVA with Dunnett’s or Sidak’s post-test was performed where indicated.

## Results

### Development of an in vitro cell engraftment model

To systematically compare cell engraftment conditions, we developed an *in vitro* engraftment model using the ALI culture system (Supplemental Figure 1). In this model, the apical surface of a well-differentiated ALI culture can be injured using a variety of methods. Recovery from injury can be tracked over time by measuring transepithelial electrical resistance (TEER). At the time of peak injury (i.e., the lowest point in TEER), exogenous cells are delivered to the apical surface of the injured ALI culture. The engrafted ALI culture is then incubated for 24h before thorough washing and downstream analysis which includes measuring percent chimerism by droplet digital PCR, evaluating CFTR ion transport in Ussing chambers, and visualizing the interactions between delivered cells and the injured airway epithelium by time-lapse confocal microscopy.

### Cells grown by the CRC method are superior for engraftment

HBECs can be collected from CF lungs by bronchiolar lavage, induced sputum sampling, or bronchoscopy which yields between 2 × 10^3^ and 2 × 10^6^ cells on average (26–28). With such meager starting materials, cell expansion on tissue culture plastic will likely be required. Several cell expansion techniques exist for airway epithelial cells including conventional culture in non-proprietary bronchial epithelial growth medium (BEGM) (29, 30) and conditionally reprogrammed cell (CRC) culture methods requiring co-culture with irradiated 3T3J2 fibroblasts and a rho kinase inhibitor (22). Previously, we have shown that HBECs grown by the CRC method have greater clonal growth capacity and competitive growth advantages compared to conventionally grown cells (31). However, the effect of CRC culture on engraftment efficiency has not been studied.

To determine if the cell expansion method affects “engraftability”, primary HBECs were expanded using BEGM or CRC methods to the first passage (P1) and transduced with an GFP-expressing adenovirus (Ad-GFP). Well-differentiated ALI cultures were preconditioned with 0.125% PDOC, a chemical injury previously used in gene and cell therapy models (11, 32–35) and tracked over time by TEER measurements. PDOC treatment diminished TEER to 1.7% of the baseline resistance in the first 5 min and remained low for the first 24 h post-injury (Figure 1A). At 24 h, BEGM P1 cells, CRC P1 cells, or a vehicle control were added to the injured ALI culture. TEER recovery was not significantly altered by the addition of BEGM P1 cells. However, CRC P1 cell addition caused a significant increase in TEER at 72h (Figure 1A). By fluorescent microscopy, we found that CRC P1 cell engraftment was nearly six-fold greater than BEGM P1 cell engraftment at 48h postinjury (i.e., 24h after cell addition) (Figure 1B; D-F). To determine if this could be explained by innate differences in cell attachment, BEGM P1 or CRC P1 cells labeled with Ad-GFP were seeded on a bare collagen-coated cell culture membrane and imaged after 24h (Figure 1C; G-H). CRC P1 cells displayed nearly four-fold greater attachment than BEGM P1 cells. From this, we concluded that cells grown by the CRC method display enhanced “engraftability” due to improved cell attachment.

**Figure 1.**
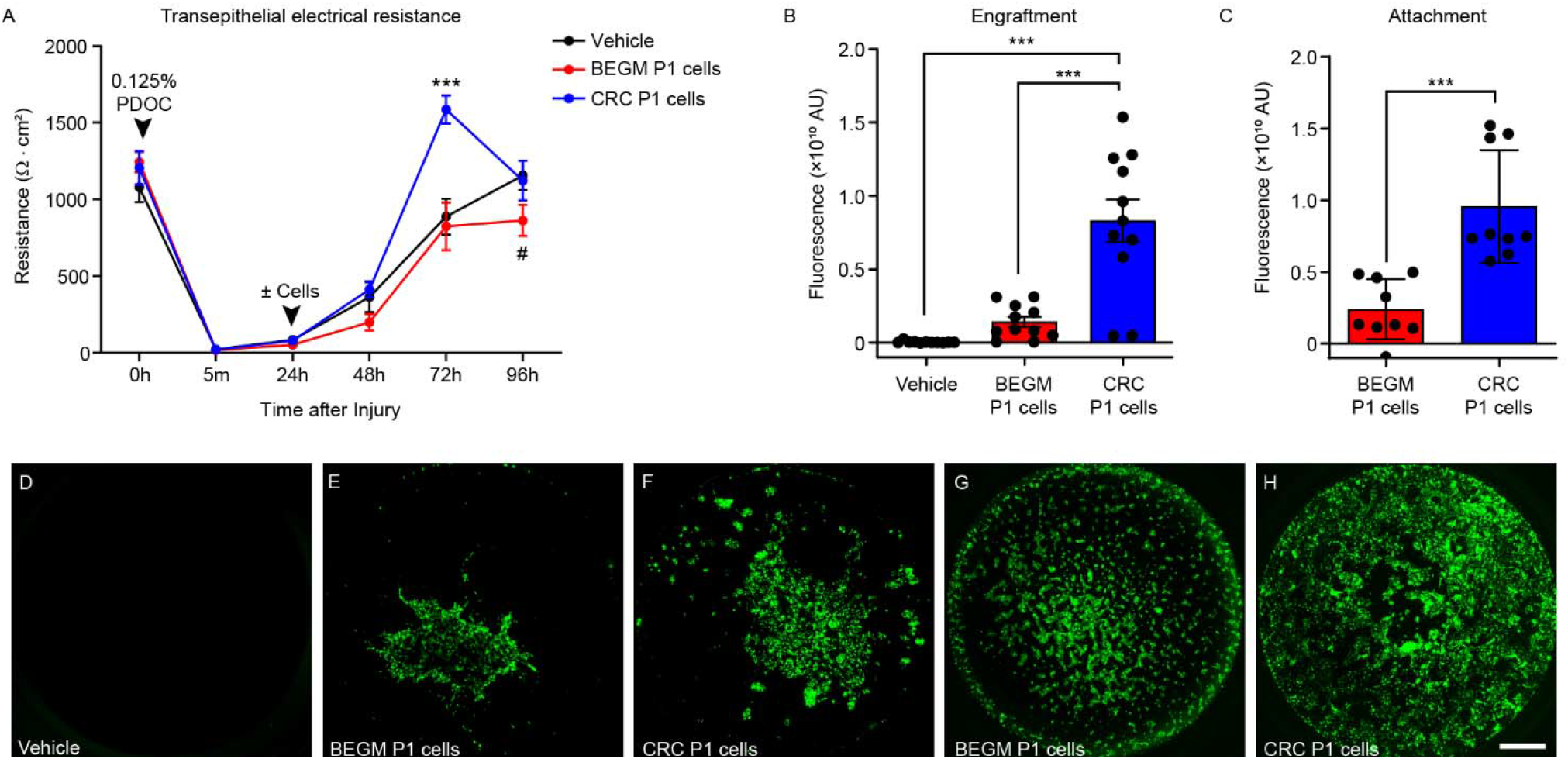
Engraftment and attachment of human bronchial epithelial cells grown in bronchial epithelial growth medium (BEGM) or by the conditionally reprogrammed cell (CRC) culture method. A) Transepithelial electrical resistance (TEER) recovery after injury with 0.125% polidocanol (PDOC) and delivery of vehicle, BEGM P1 cells, or CRC P1 cells. Two-way ANOVA with Dunnett’s multiple comparisons post-test. Vehicle vs CRC P1 cells: *** = p<0.001. Vehicle vs BEGM P1 cells: # = p<0.05. B) Quantitation of fluorescent signal expressed in arbitrary units (AU) from well-differentiated ALI cultures treated with vehicle, BEGM P1 cells, or CRC P1 cells after injury with 0.125% PDOC. One-way ANOVA with Tukey post-test. *** = p<0.001. N = 4 donors; 2-3 cultures per donor. C) Quantitation of fluorescent signal from BEGM P1 cells or CRC P1 cells seeded directly onto a collagen-coated cell culture membrane. B-C) Unpaired T-test. *** = p<0.001. N = 3 donors; 3 cultures per donor. D-F) Representative fluorescent image of well-differentiated ALI culture treated with vehicle (D), BEGM P1 cells (E), or CRC P1 cells (F) after injury with 0.125% PDOC. G-H) Representative fluorescent image of BEGM P1 cells (G) or CRC P1 cells (H) seeded directly onto a cell culture membrane. D-H) Scale bar in (H) = 1 mm.

### Characterization of sodium caprate and polidocanol injuries in vitro

Airway epithelial disruption is necessary to facilitate cell engraftment *in vivo* (35–37), however the type and extent of injury required is unknown. Using our *in vitro* engraftment model, we compared three kinds of airway injury by treating well-differentiated ALI cultures with sodium caprate (C10) to transiently disrupt tight junctions, low-dose PDOC (0.125%) to prompt luminal cell shedding, and high-dose PDOC (0.25%) to induce partial epithelial denudation and compared all injuries to a vehicle control (PBS). C10 has previously been shown to increase gene transfer (38–40) but has not been tested as a means to increase cellular uptake.

PBS-treated ALI cultures exhibited an organized, pseudostratified epithelium with abundant ciliated and goblet cells and an average epithelial height of 18.2 ± 0.1 μm (Figure 2A, E). This organized epithelium was disrupted by C10 treatment, which prompted tight junction disorder and some cell loss, reducing epithelial height to 11.6 ± 0.1 μm (Figure 2B, E). Low-dose PDOC induced luminal cell shedding, leaving only a basal cell layer intact, and yielding a short 9.7 ± 0.1 μm monolayer (Figure 2C, E). Treatment with high-dose PDOC induced more dramatic cell loss, with patches of epithelial denudation down to the basement membrane (Figure 2D, E). ALI cultures subjected to high-dose PDOC were approximately 4.2 ± 0.01 μm tall and were 42.4 ± 6.7% denuded by area (Figure 2I). All injury models reduced the number of nuclei, the ciliated cell count, and the amount of MUC5AC staining compared to the PBS-treated controls (Figure 2F-H).

**Figure 2.**
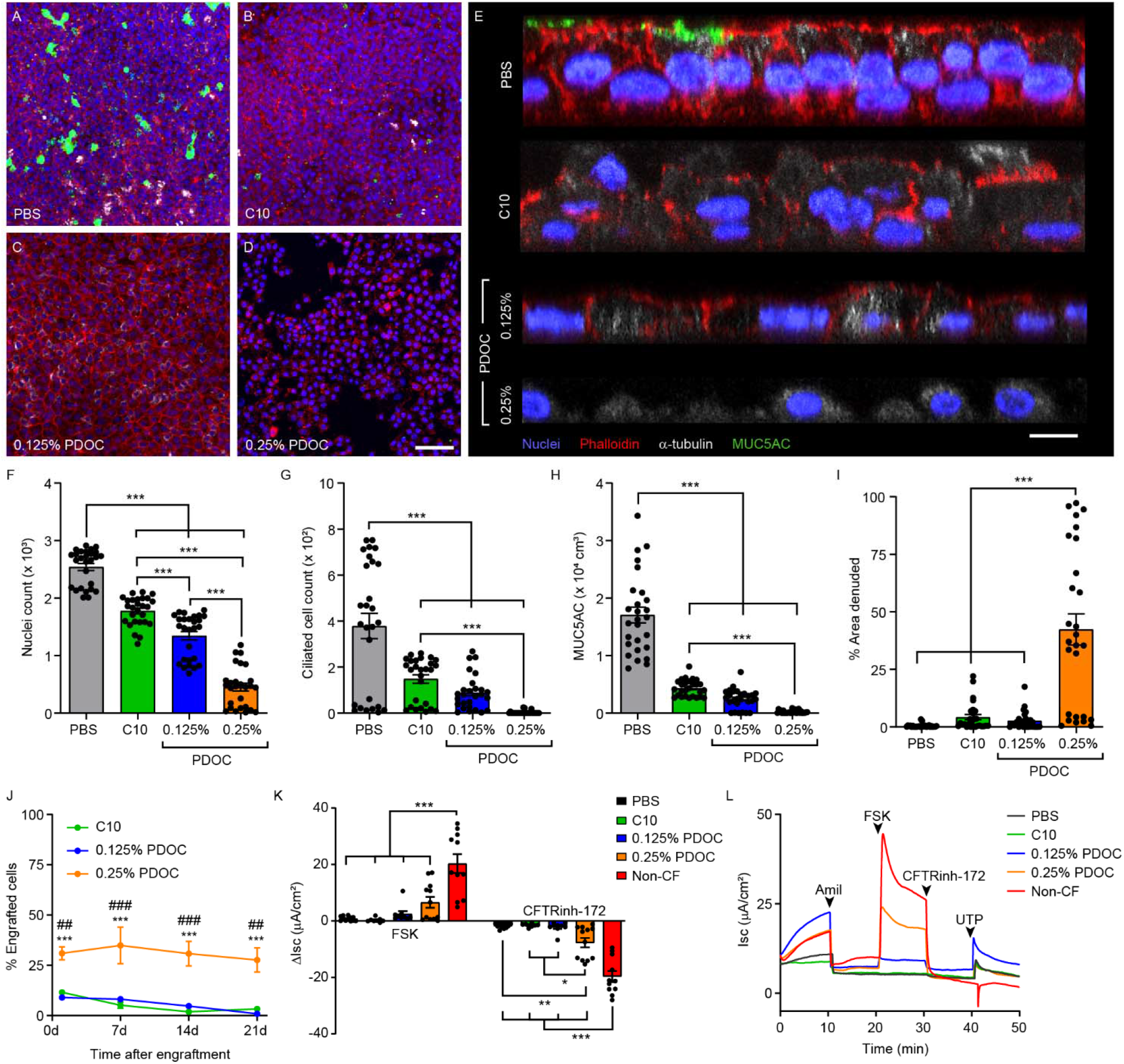
Characterization of sodium caprate and polidocanol injury of well-differentiated ALI cultures. A-D) Representative confocal microscopy image of cultures five minutes after treatment with PBS (A) or C10 (B) or 24h after treatment with 0.125% (C) or 0.25% PDOC (D). A-D) Scale bar in (D) = 50 μm. E) Orthogonal view of treated cultures. Scale bar = 10 μm. Blue = Hoechst (nuclei); red = phalloidin; white = α-Tubulin; green = MUC5AC. F-I) Quantitation of nuclei (F), ciliated cells (G), MUC5AC^+^ cells (H), and the percent of denuded area (I) per 291 × 291 μm field. One-way ANOVA with Tukey post-test. *** = p<0.001. N = 3 donors; 9 fields per donor. J) Percent of engrafted cells at 1, 7, 14, and 21 days after cell addition by droplet digital PCR. Twoway ANOVA with Dunnett’s multiple comparisons post-test. 0.25% PDOC vs 0.125% PDOC: *** = p<0.001. 0.25% PDOC vs C10: ^##^ = p<0.01, ^###^ = p<0.001. N = 3-4 donors; 2-3 cultures per donor. K) Change in short circuit current (ΔIsc) in response to Amiloride (Amil), forskolin (FSK), CFTR inhibitor-172 (CFTRinh-172), and uridine 5’-triphosphate (UTP). One-way ANOVA with Tukey post-test. * = p<0.05, ** = p<0.01, *** = p<0.001. N = 3-4 donors; 2-3 cultures per donor. L) Representative Ussing tracing of a non-CF culture and CF cultures treated with PBS, C10, 0.125% PDOC, or 0.25% PDOC and engrafted with non-CF CRC P1 cells.

### Engrafted cells persist long-term and contribute to functional ion transport in vitro

Next, we injured ALI cultures from CF donors with C10, low-dose PDOC, or high-dose PDOC and delivered non-CF cells grown by the CRC method. Engrafted cultures were lysed at 1d, 7d, 14d, or 21d post-engraftment and chimerism was analyzed by ddPCR. Twenty-four hours after cell addition, engrafted cells constituted 11.5 ± 0.9% of C10 treated cultures, but this proportion decreased at later time points, dwindling to 3.3 ± 1.0% after 21d (Figure 2J). Likewise, low-dose PDOC treated cultures contained 9.0 ± 0.7% engrafted cells at 1d post-engraftment but diminished to 0.9 ± 0.3% after 21d. By contrast, the percent of engrafted cells was three-fold higher in the high-dose PDOC treated cultures at 1d (31.0 ± 3.3%) and this proportion was maintained for at least 21d (27.7 ± 6.0%). From this, we concluded that severe injury characterized by partial epithelial denudation promotes long-term persistence of engrafted cells in the airway epithelium. Injuries that only cause tight junction disruption or luminal cell loss are insufficient.

Next, we assessed the functional contribution of the engrafted non-CF cells by measuring CFTR ion transport in Ussing chamber assays (Figure 2K-L). As expected, PBS treated CF cultures displayed no CFTR function as indicated by a negligible response to forskolin (FSK). Engrafted cultures preconditioned with C10 or low-dose PDOC also produced little to no CFTR activity. However, engrafted cultures preconditioned with high-dose PDOC yielded 20.2 ± 5.9% of wildtype CFTR function at 14d post-engraftment (Figure 2K), indicating that severe airway injury modeled by high-dose PDOC can facilitate persistent cell engraftment and that engrafted cells are capable of differentiation and reconstitution of CFTR ion transport.

### Exogenous cells expedite TEER recovery after partial epithelial denudation

To examine the dynamics of repair from partial epithelial denudation, well-differentiated ALI cultures were treated with high-dose PDOC and recovery was tracked over time with the addition of Ad-GFP labeled CRC P1 cells or a vehicle control (Figure 3A). Confocal microscopy indicated that ALI cultures were 53.4 ± 7.2% denuded 24h post-injury (Figure 3B). At 48h, vehicle-treated cultures remained 48.8 ± 7.1% denuded whereas cultures treated with CRC P1 cells were only 18.7 ± 3.7% denuded. We also tracked recovery from injury by measuring TEER (Figure 3C). TEER dropped to 1.9% of the baseline resistance within 5 min of high-dose PDOC treatment and remained low after 24h as expected. No difference was detected between treatment groups at 48h. However, TEER was significantly greater in CRC P1 treated cultures than vehicle-treated controls at both 72h and 96h. Thus, recovery from epithelial denudation is expedited by the addition of exogenous cells. There are two possible explanations for why this may be occurring: 1) exogenous cell addition could be accelerating the spread of the injured and repairing epithelium, or 2) delivered cells may be engrafting and filling in the denuded space.

**Figure 3.**
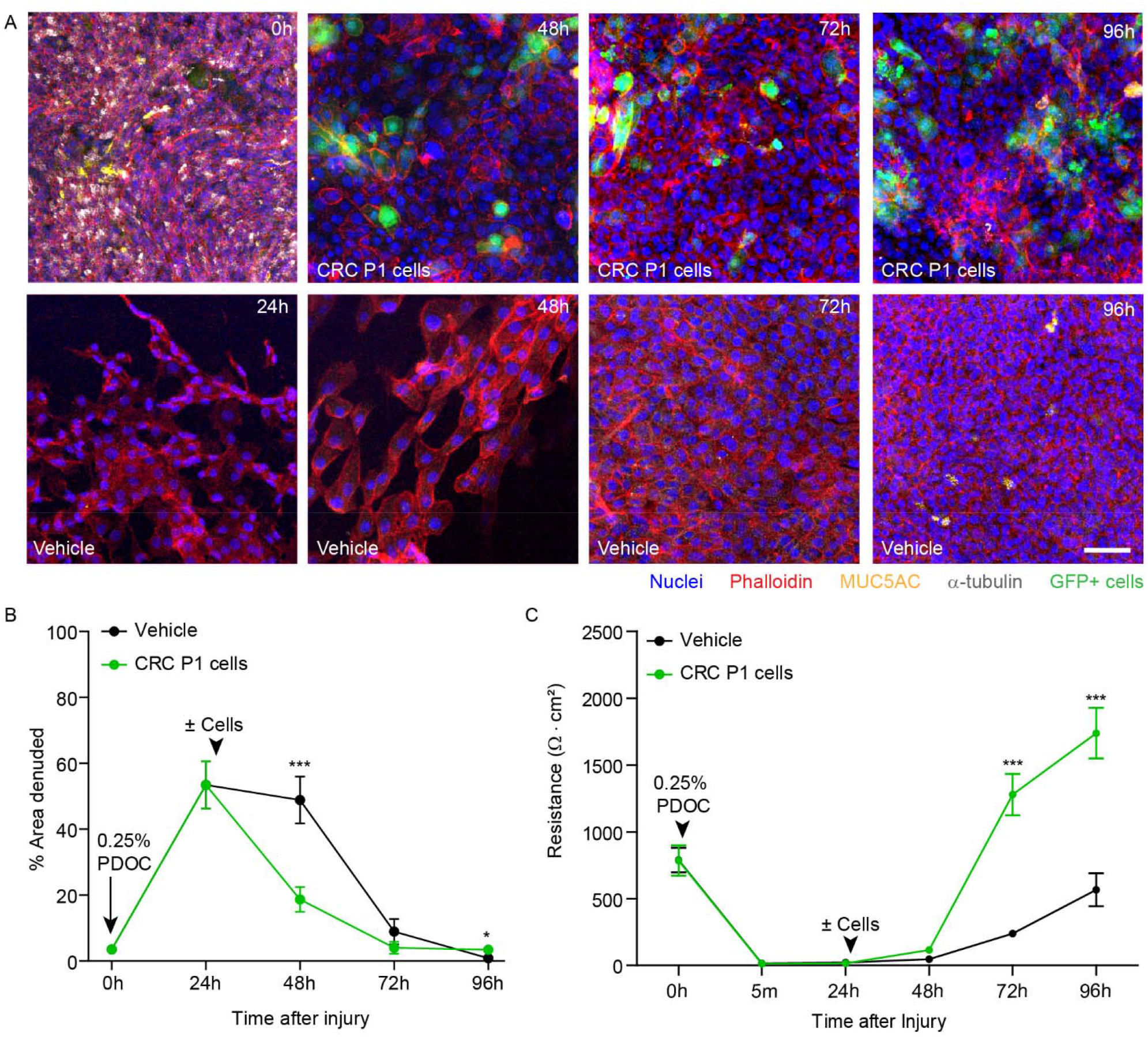
Recovery from high-dose PDOC with and without cell addition. A) Representative confocal images depicting the recovery time course from 0.25% PDOC injury followed by treatment with vehicle or GFP-labeled CRC P1 cells. Blue = Hoechst (nuclei); red = phalloidin; yellow = MUC5AC; white = α-Tubulin; green = GFP^+^ cells. Scale bar = 50 μm. B) Percent area denuded per 291 × 291 μm field after injury with 0.25% PDOC followed by treatment with vehicle or CRC P1 cells. N = 3 donors; 9 fields per donor. Unpaired T-test for each time point. * = p<0.05; *** = p<0.001. C) TEER recovery after injury with 0.25% PDOC followed by treatment with vehicle or CRC P1 cells. Unpaired T-test for each time point. *** = p<0.001. N = 3 donors; 3 cultures per donor.

To explore the first possibility, we used time-lapse confocal imaging to visualize the spread of injured and repairing airway epithelium. Well-differentiated ALI cultures were injured using high-dose PDOC and visualized with a fluorescent Calcein dye. CRC P1 cells were then stained with a mis-matched Calcein dye and added to the injured cultures as before (Figure 4A-B). Immediately following cell addition, the cultures were imaged by confocal microscopy every 20-25 min over 20h. Without cell addition, the repairing epithelium spread to cover the exposed cell culture membrane at a rate of 755.5 ± 64.8 μm^2^/min by flattening and migrating as a unified cell front. The addition of exogenous cells significantly reduced the rate of cell spread to 415.3 ± 86.6 μm^2^/min (Figure 4C-D). Conceptually, this indicates that engrafting cells likely create resistance to epithelial spread by attaching and taking up denuded space. From this, we concluded that the improved TEER recovery observed upon exogenous cell addition (Figure 3) was due to the successful incorporation and engraftment of delivered cells.

**Figure 4:**
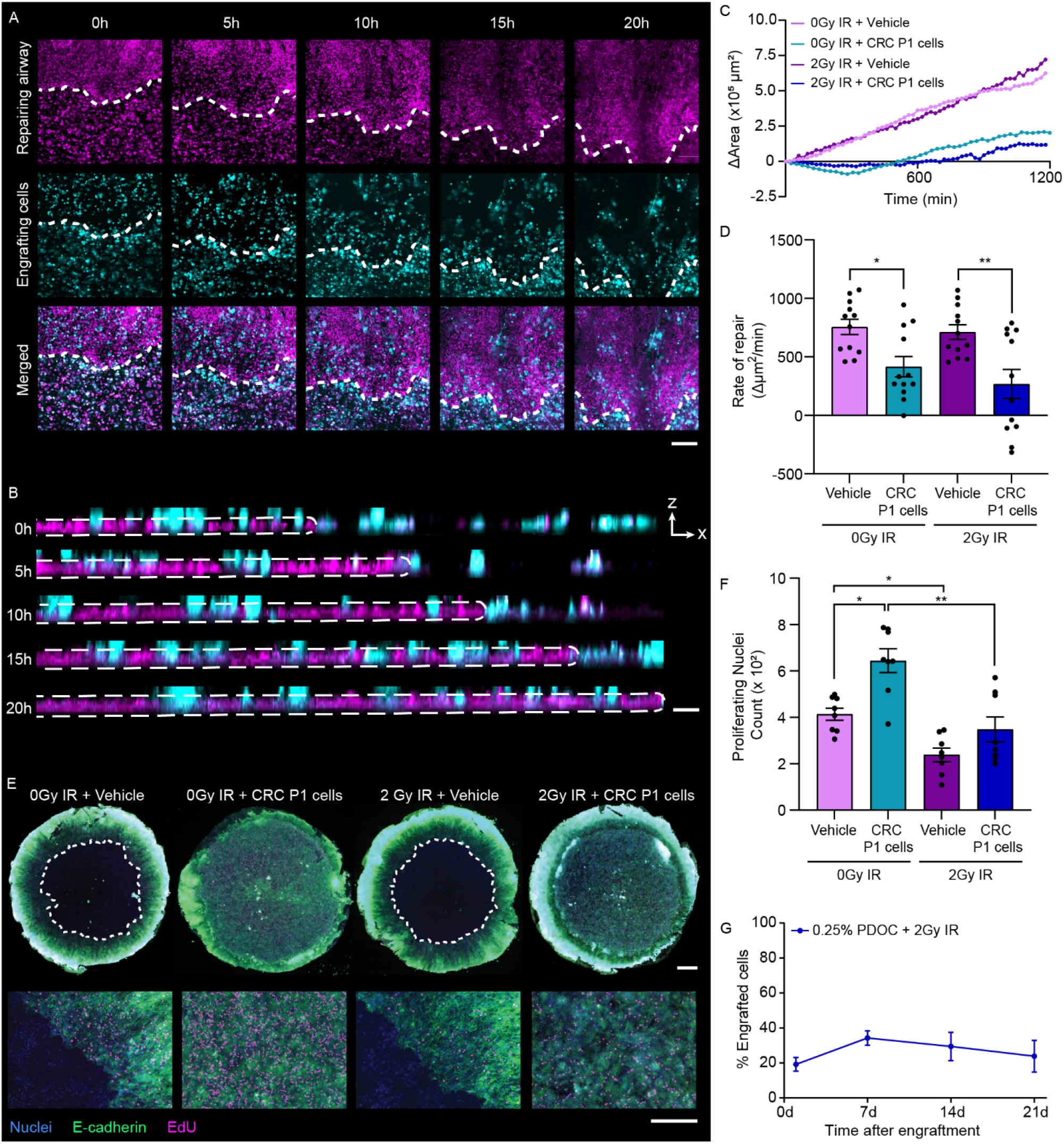
Radiation gives engrafting cells a competitive advantage by reducing the proliferation of resident stem cells. A) Representative images of a well-differentiated ALI culture (magenta) injured with 0.25% PDOC and engrafted with CRC P1 cells (cyan). Scale bar = 250 μm. B) Time course of orthogonal slices. The white dashed line marks the boundary of the repairing epithelium in A and B. Scale bar = 50 μm. C) Representative plot of the culture area covered by repairing epithelium over time in cultures injured with 0.25% PDOC and treated ± 2Gy IR and ± CRC P1 cells at 24h post-injury. D) Rate of repair (change in area over time) in cultures injured with 0.25% PDOC and treated ± 2Gy IR and ± CRC P1 cells at 24h post-injury. One-way ANOVA with Tukey post-test. * = p<0.05. ** = p<0.01. N = 2-3 donors; 1-2 cultures per donor; 4 fields per culture. E) Whole-mount immunostaining of well-differentiated ALI cultures injured with 0.25% PDOC and treated ± 2Gy IR and ± CRC P1 cells at 24h post-injury. Blue = Hoechst (nuclei); green = e-cadherin; magenta = EdU (proliferating nuclei). Top: scale bar = 1 mm. Bottom: scale bar = 250 μm. F) Quantitation of proliferating nuclei per 291 × 291 μm field. One-way ANOVA with Tukey post-test. * = p<0.05. ** = p<0.01. N = 2 cultures; 4 fields per culture. G) Percent of engrafted cells in cultures injured with 0.25% PDOC and 2Gy IR at 1, 7, 14, and 21 days after cell addition by droplet digital PCR. N = 3-5 donors; 2 cultures per donor.

### Radiation gives engrafting cells a competitive advantage by reducing the proliferation of resident stem cells

Rapid repair after airway injury is vital to reestablish barrier function in the lungs. However, this regenerative process creates competition with delivered cells which need physical space to attach. Previous work by Rosen and colleagues demonstrated that total body irradiation (IR) improved cell engraftment in a murine naphthalene injury model by giving delivered cells a competitive advantage (36). Thus, we hypothesized that IR would shift the competition dynamics in favor of engrafted cells in our ALI model system. ALI cultures injured with high-dose PDOC were further conditioned with 2 gray (Gy) of IR before CRC P1 addition and time-lapse confocal imaging. Radiation did not alter the rate of epithelial repair regardless of whether or not cells were added (Figure 4C-D).

Next, we evaluated cell proliferation by staining engrafted ALI cultures with Click-iT EdU (Figure 4E). The rate of proliferation was significantly higher in ALI cultures with added CRC P1 cells (Figure 4F). However, we could not distinguish between proliferation of resident stem cells of the repairing epithelium and engrafting cells by this method. By contrast, IR significantly reduced the rate of proliferation ± delivered cells (Figure 4F). Initial cell engraftment at 1d appeared to be similar with or without IR (Figure 2J; Figure 4G). However, the percent of engrafted cells increased by 7d in cultures treated with high-dose PDOC and 2Gy IR, indicating an expansion of the engrafted cell population that was not seen in PDOC preconditioning alone (Figure 2J). From this, we concluded that radiation gives engrafting cells a competitive edge by diminishing resident stem cell proliferation.

### Viral injury promotes cell engraftment into the airway epithelium

We demonstrated that cell engraftment into the airway epithelium was best facilitated by injuries that caused partial epithelial denudation. However, intentionally injuring lungs to this extent would be dangerous. Viral infections are known to cause airway damage reminiscent of the high-dose PDOC injury, with epithelial sloughing and patches of denudation (15, 16). Thus, we hypothesized that viral damage to the airway epithelium could serve as a naturally occurring injury regimen to prime the airways for cell engraftment. One potential concern about delivering cells to virus-infected airways is that delivered cells might also become infected, preventing effective engraftment. To address this, we hypothesized that viral resistance could be promoted in delivered cells by pretreating the engrafting cells with interferon-λ (IFN-λ).

Using our *in vitro* engraftment model, we inoculated well-differentiated CF ALI cultures with either influenza-A or vehicle (mock inoculation). At the peak of injury (5d post-infection), cultures were treated with vehicle control or 100 μg/ml ribavirin, an antiviral nucleoside, to halt the spread of virus. Non-CF CRC P1 cells were pretreated with or without 1.2 ng/ml IFN-λ to promote viral-resistance and were engrafted into CF ALI cultures at 6d post-infection (Supplemental Figure 2). We first examined the morphology of mock inoculated or influenza-A infected ALI cultures treated ± ribavirin at day 5 (Figure 5A). In the absence of virus, ribavirin-treated cultures remained confluent. Similarly, cultures inoculated with virus only remained confluent. However, the combination of influenza-A and ribavirin led to partial epithelial denudation reminiscent of high-dose PDOC.

**Figure 5.**
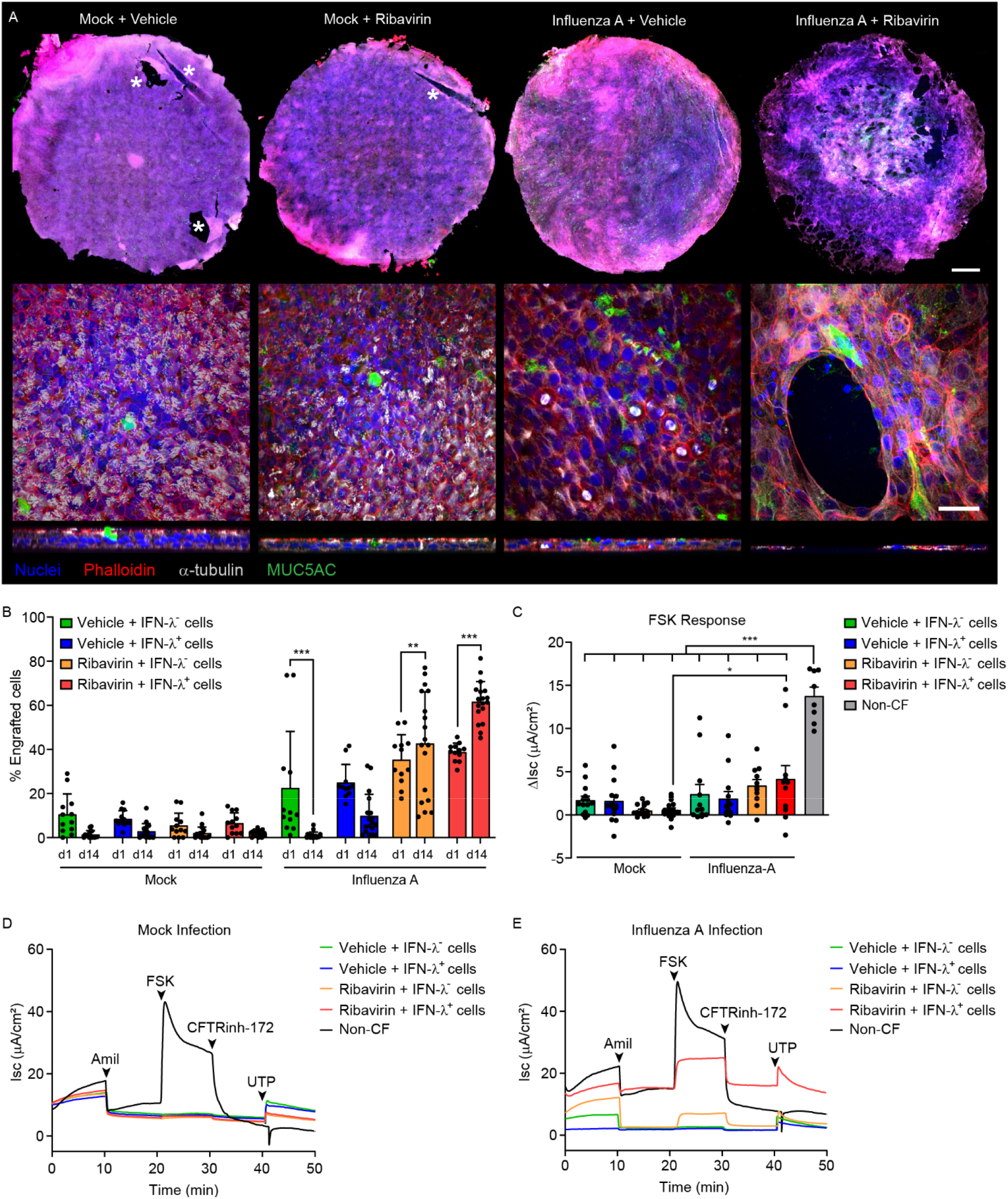
Characterization of engraftment following viral preconditioning. A) Representative confocal images of mock inoculated or influenza-A infected cultures treated ± ribavirin on day 5 post-infection (see Supplemental Figure 1). Asterisk (*) marks regions where the cell culture membrane was damaged by forceps in the immunostaining process. Blue = Hoechst (nuclei); red = phalloidin; white = α-Tubulin; green = MUC5AC. Top: scale bar = 1 mm. Bottom: scale bar = 50 μm. B) Percent of non-CF cells at 1d and 14d by ddPCR in mock and influenza-A inoculated cultures treated ± ribavirin and engrafted with CRC P1 cells pretreated ± IFN-λ. Two-way ANOVA with Sidak’s multiple comparisons post-test between timepoints (** = p<0.01; *** = p<0.001). Two-way ANOVA with Tukey’s post-test between treatments reported in Table 1. N = 3 donors; 3-4 cultures per donor. C) ΔIsc in FSK response. One-way ANOVA with Tukey’s post-test between treatments (* = p<0.05; *** = p<0.001). D-E) Representative Ussing tracings of cultures inoculated with mock (D) or influenza-A (E) and treated ± ribavirin and with CRC P1 cells pretreated ± IFN-λ. N = 3-5 donors; 1-3 cultures per donor.

Next, we evaluated chimerism at 1d and 14d post-engraftment by ddPCR (Figure 5B). All vehicle-treated cultures exhibited low levels of engraftment, reaching only 5.5 – 10.6% engrafted cells at 1d and diminishing to 1.4 – 2.9% engrafted cells by 14d. In contrast, influenza-treated cultures had much higher levels of initial engraftment, with 22.5 – 33.8% engrafted cells at 1d. The percent of engrafted cells further increased over 14d to 42.7 – 62.6% in ribavirin-treated, influenza-conditioned cultures, but without ribavirin, the percent of engrafted cells decreased over time, possibly due to unchecked viral replication and toxicity. Statistically, the biggest differences in cell engraftment were seen in comparisons between mock and influenza-A groups, followed by comparisons between ± ribavirin groups (Table 1). Finally, we evaluated the degree of CFTR functional restoration by engrafted non-CF cells in Ussing chamber assays (Figure 5C-E). Influenza-A infection paired with ribavirin treatment promoted greater CFTR ion transport function than other treatment groups, promoting 30.2% wildtype functional recovery on average (Figure 5C). From this, we concluded that viral damage to the airway epithelium provides sufficient preconditioning to facilitate effective long-term cell engraftment.

**Table 1.** Statistical analysis of ddPCR measurement of cell engraftment at 1d and 14d in mock and influenza-A inoculated cultures treated ± ribavirin and engrafted with CRC P1 cells pretreated ± IFN-λ. Results from a twoway ANOVA with Tukey post-test between treatment groups. N = 3 donors; 3-4 cultures per donor.

| **Group 1** | **vs** | **Group 2** | **P-Value** |
| --- | --- | --- | --- |
| **Mock** + Vehicle + IFN-$\lambda^-$ cells | vs | **Influenza-A** + Vehicle + IFN-$\lambda^-$ cells | P<0.001 |
| **Mock** + Ribavirin + IFN-$\lambda^-$ cells | vs | **Influenza-A** + Ribavirin + IFN-$\lambda^-$ cells | P<0.001 |
| **Mock** + Ribavirin + IFN-$\lambda^+$ cells | vs | **Influenza-A** + Ribavirin + IFN-$\lambda^+$ cells | P<0.001 |
| Influenza-A + **Vehicle** + IFN-$\lambda^-$ cells | vs | Influenza-A + **Ribavirin** + IFN-$\lambda^-$ cells | P<0.001 |
| Influenza-A + **Vehicle** + IFN-$\lambda^+$ cells | vs | Influenza-A + **Ribavirin** + IFN-$\lambda^+$ cells | P<0.001 |
| Influenza-A + Ribavirin + **IFN-$\lambda^-$ cells** | vs | Influenza-A + Ribavirin + **IFN-$\lambda^+$ cells** | P<0.01 |

## Discussion

Cell engraftment into the airway epithelium is a monumental barrier. Previous studies in animal models have achieved only 1 – 4% engraftment in the upper airways (10, 41, 42). Here, we introduce an *in vitro* engraftment model and demonstrate its utility to optimize cell engraftment strategies. With this model, we systematically compared engrafting cell populations and injury methods, tracked the behavior of engrafting cells, and evaluated a naturally occurring airway injury for its ability to facilitate cell engraftment.

Low attachment of delivered cells significantly limits the efficacy of cell therapy and is the primary reason that ample cellular resources are required (11). As such, identifying cells with enhanced “engraftability” could prove invaluable. We found that cells grown by the CRC method are superior for attachment and engraftment into the airway epithelium. This may be due to increased expression of adhesion proteins like integrin α6 and β1 (22, 29, 43) or increased competitiveness with endogenous stem cells as previously suggested (31). The CRC method offers numerous advantages towards making cell therapy possible. First, CRC expansion permits extensive growth of patient-derived cells with more population doublings per passage than conventional culture methods (44, 45). Second, cells grown by the CRC method are efficiently transduced by viral vectors (46). Studies here were performed by engrafting non-CF cells into CF ALI cultures, but clinically, the goal would be to collect cells from a person with CF, correct the CFTR gene *ex vivo,* and engraft back into the lungs (i.e., autologous cell therapy). Accordingly, a cell population that could be efficiently transduced with a CFTR-correcting vector would be highly advantageous.

Here, we demonstrate that effective engraftment requires partial epithelial denudation to deplete resident stem cells and create space for attachment of delivered cells. Our *in vitro* findings align with *in vivo* work (35, 37) and may explain why milder injuries in other *in vivo* studies fail to promote long-term cell retention (10, 12). Some studies have demonstrated the ability of engrafted cells to differentiate into secretory and ciliated cell types (47–49). Yet, to our knowledge, we are the first to show that engrafted cells can restore functional ion transport. In a foundational study of CFTR gene correction, Zhang et al., found that 25% correction of airway luminal cells fully restored CFTR function (50). However, we found that when the final CF culture was composed of 28% engrafted non-CF cells, CFTR function was corrected to ~20% of wildtype function. One possibility is that engrafted cells in our model occupy both basal and luminal cell compartments, and that the percent of luminal cells corrected is lower than 25%. Basal cell correction would be advantageous for the long-term efficacy of cell therapy.

We also found that the addition of exogenous cells accelerated repair from partial epithelial denudation. Intriguingly, this was not attributed to an increase in the rate of cell spreading to cover denuded space, but by exogenous cell attachment and incorporation into the epithelium. In a seminal study of competition, Rosen et al., demonstrated that total body irradiation following naphthalene injury in a mouse model significantly reduced resident stem cell proliferation and granted a competitive advantage to the engrafting cell population (36). Likewise, we observed a decrease in endogenous proliferation following IR and an expansion of engrafted cells over time.

Finally, we demonstrate *in vitro* that viral injury to the airway epithelium is sufficient preconditioning for effective cell engraftment. Exploiting the post-viral milieu may abrogate the need to intentionally injure the lungs to facilitate cell therapy for CF. Though influenza-A was used in this study, our findings may extend to other viruses that commonly affect people with CF such as respiratory syncytial virus (RSV). The absence of an immune system *in vitro* is a shortcoming. Thus, we used ribavirin to halt viral spread. In ribavirin treated cultures, the percent of engrafted cells drastically increased over time. This could be explained by the continued death of virally infected CF cells, a competitive advantage of engrafting cells due to their viral resistance, or both. Ultimately, *in vivo* studies to determine how the immune system’s response to viral infection affects engraftment efficiency will be critical.

The concept of delivering cells primed for viral resistance to infected airways has therapeutic potential beyond CF. Globally, severe viral infection represents a major health burden with ~60,000 deaths per year caused by respiratory syncytial virus (RSV) (51), 300,000-650,000 deaths per year caused by influenza-A and-B (52), and ~15 million deaths in two years since the emergence of the severe acute respiratory syndrome coronavirus-2 (SARS-CoV-2) in 2019 (53). Unlike cell therapy for CF where cell engraftment is key, the primary goal of cell therapy for severe viral infection would be to promote lung repair. Preliminary clinical results of mesenchymal stem cell (MSC) therapy for bronchiectasis (54), chronic obstructive pulmonary disease (COPD)/emphysema (55–57), and acute respiratory distress syndrome (ARDS) (58, 59) indicate that delivered cells can promote lung repair. Cell therapy for severe viral infection is a logical extension of these clinical studies. However, recent attempts to deliver MSCs to influenza-infected mice were unsuccessful with no improvements in markers of lung injury (60, 61). One possible explanation is that the delivered MSCs became infected with virus, a known phenomenon (62). Extending our work to these findings, we suggest that delivering cells primed for viral resistance may enhance the efficacy of cell therapy for severe viral infection.

Overall, these findings demonstrate that cell engraftment into the airway epithelium is immensely challenging, but possible. Severe injury causing partial epithelial denudation will be required unless creative methods to further enhance cell engraftment are developed. The extent of injury required may appear discouraging, however the potential for long-term functional CFTR restoration is compelling. Ultimately, more work is required to develop cell therapy into a safe and effective treatment for cystic fibrosis.

## Acknowledgements

We gratefully acknowledge the UNC Marsico Lung Institute Tissue Procurement and Cell Culture Core.

## Notes

Supported by CFF grants RANDELXX015, RANDELXX017, RANDEL20XX2, and BOUCHE19R0 and NIH grant P30DK065988, 1T32GM133364, and 1F31HL158197.

### Competing Interest Statement

The authors have declared no competing interest.

