## Supplement for "Viral Airway Injury Promotes Cell Engraftment in an *In Vitro* Model of Cystic Fibrosis Cell Therapy"

**Running Title:** Virus Injury Promotes Engraftment *In Vitro*

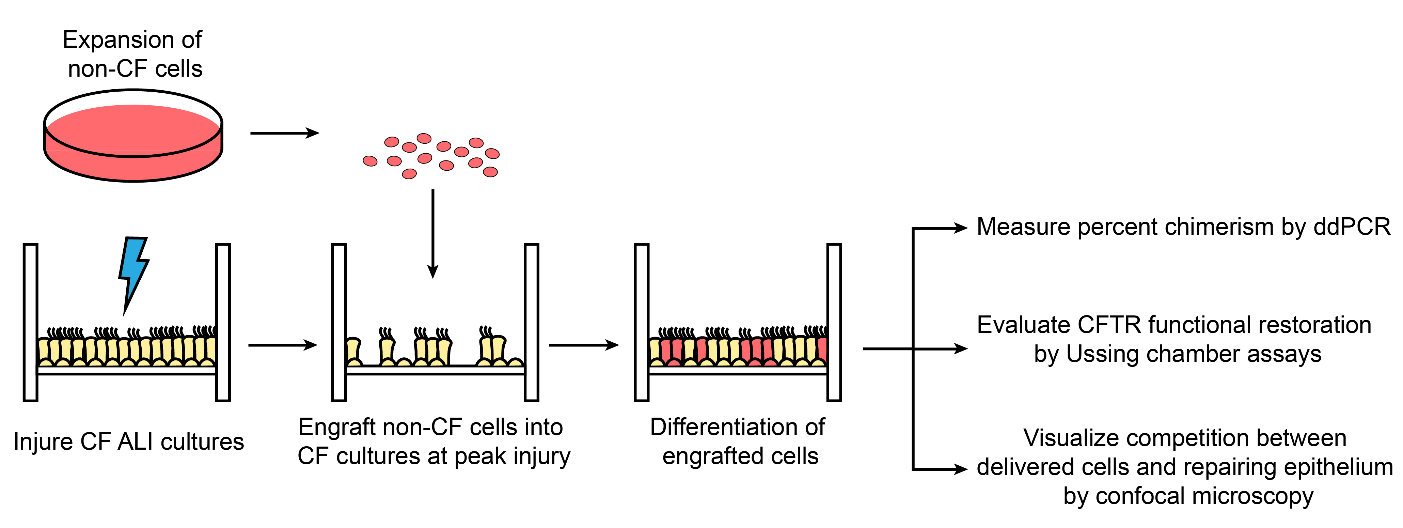

*Supplemental Figure 1. In vitro model of cell engraftment.* Well-differentiated ALI cultures from CF donors were injured using a variety of methods. At the time of peak injury, non-CF cells were delivered to the apical surface of the injured ALI culture. The engrafted ALI cell culture was then incubated for 24h before thorough washing and downstream analysis which included assessing percent chimerism by ddPCR, evaluating CFTR ion transport in Ussing chamber assays, and examining competition dynamics between delivered cells and the repairing epithelium via confocal microscopy.

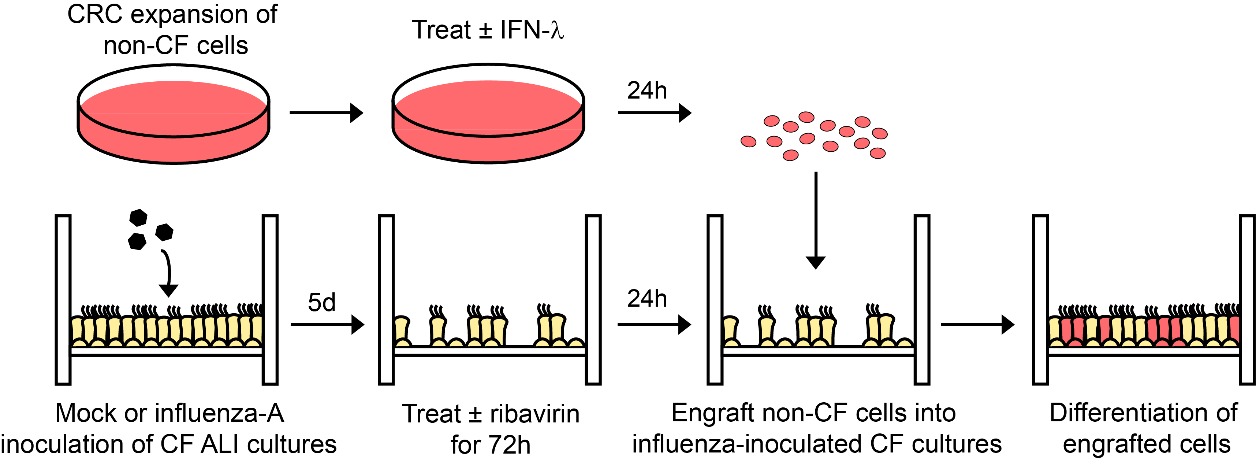

*Supplementary Figure 2. Viral preconditioning for cell engraftment*. Non-CF CRC P1 cells were treated ± IFN-λ for 24h. Concurrently, well-differentiated CF ALI cultures were inoculated with mock or influenza-A. Beginning d5 post-infection, ALI cultures were treated ± ribavirin for 72h. Twenty-four hours after ribavirin treatment began, CRC P1 cells were delivered to CF cultures. Engrafted cultures were analyzed 1d and 14d post-engraftment. Experiments were performed ± influenza-A, ± ribavirin, and ± IFN-λ pretreated cells.
